# Frameshift and frame-preserving mutations in zebrafish *presenilin 2* affect different cellular functions in young adult brains

**DOI:** 10.1101/2020.11.21.392761

**Authors:** Karissa Barthelson, Stephen Martin Pederson, Morgan Newman, Haowei Jiang, Michael Lardelli

## Abstract

**Background:** Mutations in *PRESENILIN 2* (*PSEN2*) cause early disease onset familial Alzheimer’s disease (EOfAD) but their mode of action remains elusive. One consistent observation for all *PRESENILIN* gene mutations causing EOfAD is that a transcript is produced with a reading frame terminated by the normal stop codon – the “reading frame preservation rule”. Mutations that do not obey this rule do not cause the disease. The reasons for this are debated.

**Methods:** A frameshift mutation (*psen2^N140fs^*) and a reading frame-preserving mutation (*psen2^T141_L142delinsMISLISV^*) were previously isolated during genome editing directed at the N140 codon of zebrafish *psen2* (equivalent to N141 of human *PSEN2*). We mated a pair of fish heterozygous for each mutation to generate a family of siblings including wild type and heterozygous mutant genotypes. Transcriptomes from young adult (6 months) brains of these genotypes were analysed. Bioinformatics techniques were used to predict cellular functions affected by heterozygosity for each mutation.

**Results:** The reading frame preserving mutation uniquely caused subtle, but statistically significant, changes to expression of genes involved in oxidative phosphorylation, long term potentiation and the cell cycle. The frameshift mutation uniquely affected genes involved in Notch and MAPK signalling, extracellular matrix receptor interactions and focal adhesion. Both mutations affected ribosomal protein gene expression but in opposite directions.

**Conclusion:** A frameshift and frame-preserving mutation at the same position in zebrafish *psen2* cause discrete effects. Changes in oxidative phosphorylation, long term potentiation and the cell cycle may promote EOfAD pathogenesis in humans.

## Introduction

Alzheimer’s disease (AD) is a progressive neurodegenerative disorder which develops silently over decades. The pathological hallmarks of AD include the presence of extracellular senile neuritic plaques consisting, primarily, of amyloid β (Aβ) peptides, intracellular neurofibrillary tangles (primarily consisting of hyperphosphorylated tau proteins), and progressive neuronal loss (reviewed in [1]). The majority of therapeutics for AD are aimed at reducing Aβ levels (reviewed in [2]). However, all compounds to date show little or no effect on cognitive symptoms (reviewed in [3, 4]). This likely reflects our ignorance of the pathogenic mechanism underlying AD. Also, once cognitive changes occur, damage to the brain is considerable and may be irreversible. A comprehensive understanding of the early molecular changes/stresses occurring many years before disease onset is required to develop preventative treatments to reduce the prevalence of AD.

The majority of AD cases arise sporadically with an age of onset after 65 years (late onset AD, LOAD). Genome-wide association studies (GWAS) have identified at least 20 genes associated with increased risk for LOAD, with the most influential being the *APOLIPOPROTEIN E* (*APOE*) locus [5, 6]. Rare, familial forms of AD also exist. Early onset familial AD (EOfAD) cases have an age of onset before 65 years, and arise due to mutations in the *PRESENILIN* genes (*PSEN1* and *PSEN2*), and the genes *AMYLOID B A4 PRECURSOR PROTEIN* (*AβPP*) and *SORTILIN-RELATED RECEPTOR 1* (*SORL1*) (reviewed in [7–9]).

*PSEN2* is the less commonly mutated *PSEN* gene in EOfAD. To date, only 13 pathogenic mutations have been described in *PSEN2* compared to over 185 pathogenic mutations in the homologous gene *PSEN1* [10]. The first *PSEN2* EOfAD mutation characterised was N141I, found in Volga German families and affecting the second transmembrane domain of the PSEN2 protein [11]. We previously attempted to introduce this mutation into zebrafish by genome editing. While we were not successful at introducing an exact equivalent of the human N141I mutation into zebrafish, we did isolate a reading frame-preserving mutation at this position: *psen2^T141_L142delinsMISLISV^* [12]. This mutation alters two codons and inserts five additional novel codons and was predicted to largely preserve the transmembrane structure of the resultant protein. We also isolated a frameshift mutation at the same position: *psen2^N140fs^*. This is a deletion of 7 nucleotides resulting in a premature stop codon at novel codon position 142 [12]. Since all EOfAD mutations in the *PSENs* follow a “reading frame preservation rule” [13], *psen2^T141_L142delinsMISLISV^* models such a mutation in *PSEN2*. In contrast, the *psen2^N140fs^* allele is predicted to express a truncated protein, is not EOfAD-like, and so can act as a negative control to investigate the differences between reading frame-preserving and -destructive mutations of *PSEN2*.

The overall goal of the Alzheimer’s Disease Genetics Laboratory has been to identify the cellular processes affected in common by EOfAD mutations in different genes in young adult, heterozygous mutant brains (i.e. to establish an early EOfAD brain transcriptomic signature). We exploit zebrafish as a model organism for this work as the EOfAD genes are conserved in zebrafish, and large families containing both EOfAD-like mutant fish and their wild type siblings can be generated from single mating events between single pairs of fish (reviewed in [14]). Analysing the transcriptomes of sibling fish raised in the same environment (aquaria within a single recirculated water system) reduces genetic and environmental sources of variation, (noise) and allows subtle effects of mutations to be identified. After introducing EOfAD-like mutations into the endogenous, zebrafish orthologues of EOfAD genes (i.e. knock-in models), we analyse their effects on brain transcriptomes in heterozygous fish to mimic the genetic state of such mutations in the human disease. We term this experimental strategy <u>Be</u>tween <u>S</u>ibling <u>T</u>ranscriptome (BeST) analysis. We have previously performed BeST analyses for EOfAD-like mutations in *psen1* [15] and *sorl1* [16, 17] and for a complex (but possibly EOfAD-like) mutation in *psen2*: *psen2^S4Ter^* [18].

In the work described in this report, we performed a BeST analysis using a family of sibling fish generated by mating two fish with genotypes *psen2^T141_L142delinsMISLISV^*/+ and *psen2^N140fs^*/+. The family included fish heterozygous for the mutations *psen2^T141_L142delinsMISLISV^* (for simplicity, hereafter referred to as “EOfAD-like”) and *psen2^N140fs^* (for simplicity, hereafter referred to as “FS”) and their wild type siblings, all raised together in the same tank until 6 months of age. This strategy allowed direct comparisons of the brain transcriptomes from the two *psen2* mutant genotypes to their wild type siblings while reducing confounding effects from genetic and environmental variation. Despite this, considerable gene expression variation (noise) is still observed between individuals and the effects of the mutations are subtle so that few genes are identified initially as differentially expressed. However, when analysed at the pathway level, heterozygosity for each mutation shows distinct transcriptomic effects. The sets of genes involved in oxidative phosphorylation, longterm potentiation, and the cell cycle are only altered significantly in the EOfAD-like brains, while the FS mutant brains shows apparent changes in Notch and MAPK signalling, focal adhesion, and ECM receptor interaction. Both mutation states affected genes encoding ribosomal subunits but the effect of each mutation was in opposite directions. Our results are consistent with the concept that only dominant, reading frame-preserving mutations of the *PRESENILIN* genes cause EOfAD and support that effects on oxidative phosphorylation may be a common signature of such mutations.

## Methods

### Zebrafish lines and ethics statement

Generation of the *psen2^T141_L142delinsMISLISV^* and *psen2^N140fs^* lines of zebrafish used in this study is described in [12]. All experiments involving zebrafish were conducted under the auspices of the Animal Ethics Committee of the University of Adelaide, permit numbers S-2017-089 and S-2017-073.

### Breeding strategy

Two fish heterozygous for these mutations were mated to generate a family of sibling fish with *psen2* genotypes EOfAD-like/+, FS/+, EOfAD-like/FS (transheterozygous) or wild type (**Figure 1A**). This family was raised together in the same tank until 6 months of age, at which time 24 fish were randomly selected and sacrificed in a loose ice slurry (to allow for n = 5 of each genotype in the RNA-seq analysis). The heads of these fish were removed and stored in 600 μL of RNAlater solution, while their tails were removed for genomic DNA extraction and genotyping by polymerase chain reactions (PCRs).

**Figure 1:**
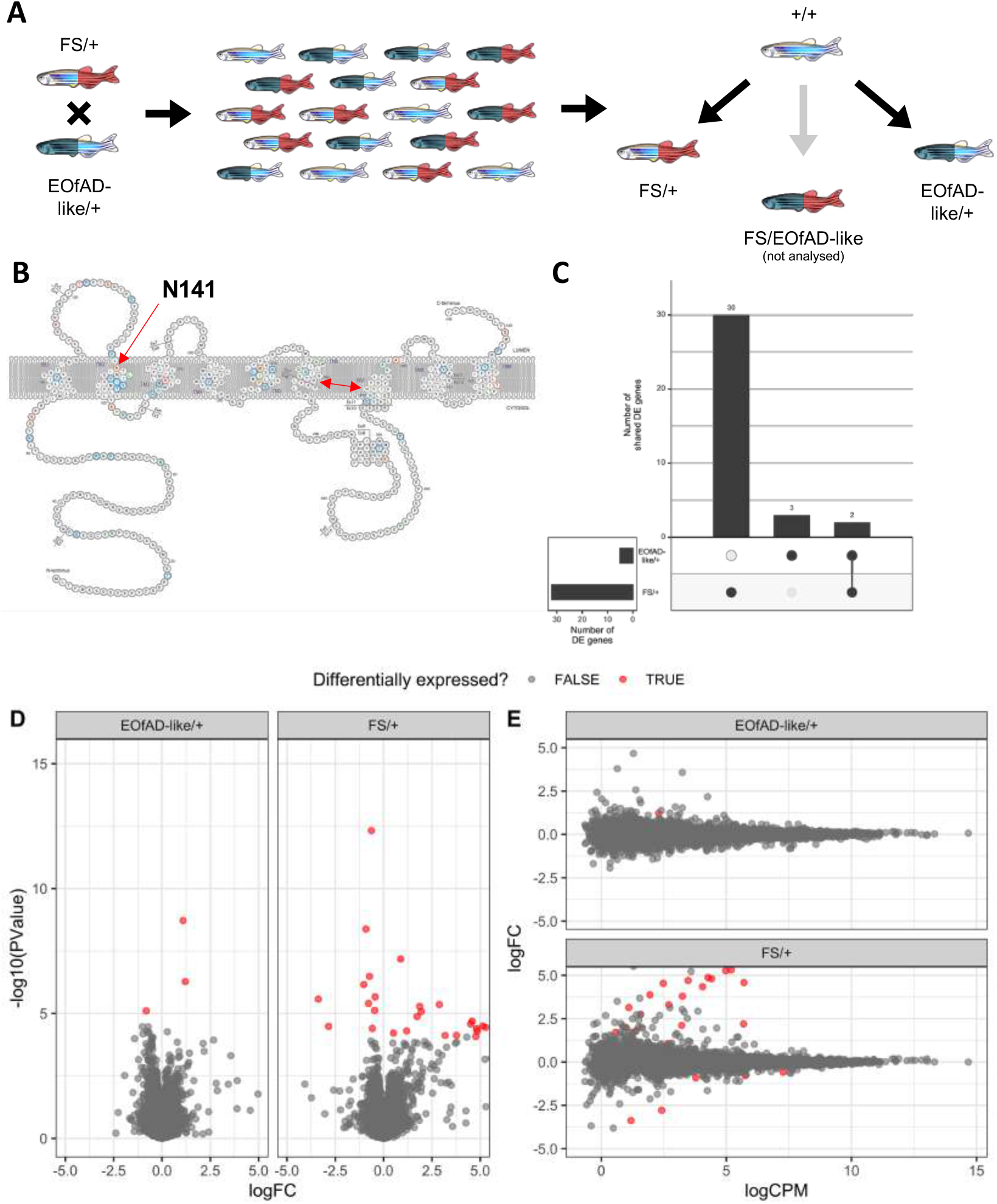
**A)** Mating strategy and experimental design. **B)** Schematic of the human PSEN2 protein (adapted from https://www.alzforum.org/mutations/psen-2 with permission from FBRI LLC (Copyright © 1996– 2020 FBRI LLC. All Rights Reserved. Version 2.7 – 2020)). Amino acid residues are color-coded to indicate whether substitutions have been observed to be pathogenic (orange), non-pathogenic (green) or of uncertain pathogenicity (blue). The site of the human EOfAD mutation N141l, in the second transmembrane domain (TMD) is indicated by the red single-headed arrow. The aspartate residues critical for γ-secretase catalysis are indicated by a red double-headed arrow. **C)** Upset plot indicating the number of differentially expressed (DE) genes in each comparison of the *psen2* mutant genotypes to wild type noting that only 2 genes appear to be DE in common between both comparisons. **D)** Mean-difference (MD) plot and **E)** volcano plot of DE genes in each comparison. Note that the logFC axis limits in D) and E) are constrained to −5 and 5 for visualisation purposes.

### PCR genotyping

Allele-specific genotyping PCRs were performed on genomic DNA extracted from fin biopsies as described in [17]. The allele-specific primer sequences can be found in **Table 1**. Primers were synthesised by Sigma Aldrich (St. Louis, MO, USA).

**Table 1:**
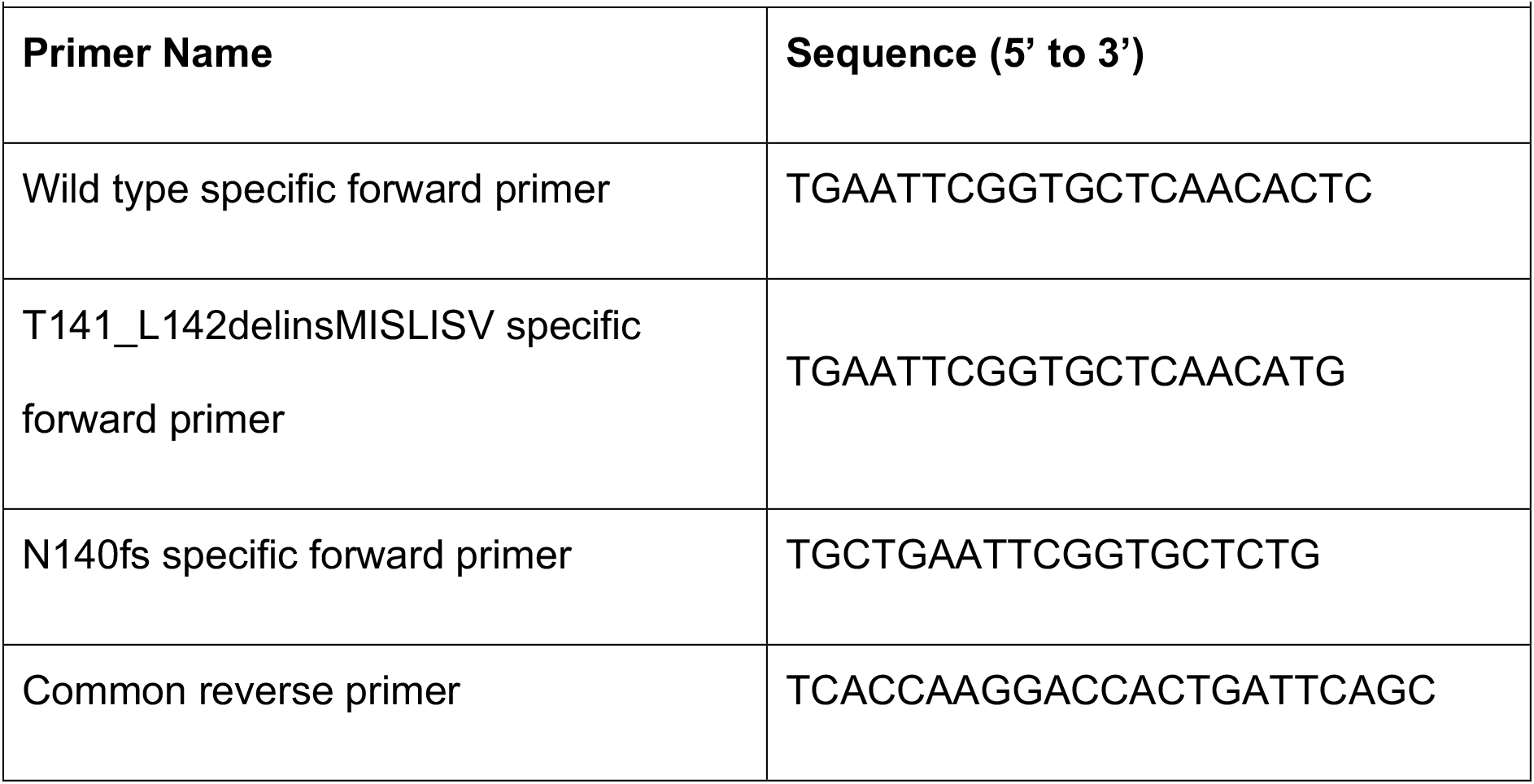
Genotyping primer sequences.

### RNA-seq data generation and analysis

Preparation of RNA for RNA-seq and the subsequent analysis of the data was performed mostly as in [17]. Briefly, brains of n = 5 fish of wild type, EOfAD-like/+ and FS/+ genotypes (each group contained 3 females and 2 males) were carefully removed from the heads preserved RNAlater Stabilization Solution (Invitrogen, Carlsbad, CA, USA), followed by total RNA extraction using the mirVana™ miRNA Isolation Kit (Ambion, Austin, TX, USA) and DNase treatment using the DNA-free™ Kit (Ambion). Total RNA was then delivered to the Genomics service at the South Australian Health and Medical Research Institute (SAHMRI, Adelaide, AUS) for polyA+, stranded library preparation and RNA sequencing using the Illumina Nextseq platform.

Processing of the demultiplexed fastq files provided by SAHMRI (75 bp single-end reads) was performed as in [17]. However, the pseudo-alignment using *kallisto* (v0.43.1) [19] was performed using a modified version of the zebrafish transcriptome (Ensembl release 94) which additionally included the two novel mutant *psen2* transcript sequences.

Transcript counts as estimated by *kallisto* were imported into *R* [20] using *tximport* [21], summing the transcript-level counts to gene-level counts. We also imported the transcript-level counts using the *catchKallisto* function from the package *edgeR* [22] to assess simply the allele-specific expression of the different *psen2* transcripts.

We removed genes considered as undetectable (less than one count per million in at least five of the samples) from downstream analysis, leaving library sizes ranging between 17 million and 27 million reads. We normalised for these differences in library sizes using the trimmed mean of M-values (TMM) method [23], followed by removal of one factor of unwanted variation using the *RUVg* method *RUVseq* [24] as described in [17].

Differential gene expression analysis was performed using a generalised linear model (GLM) and likelihood ratio test using *edgeR* [22], specifying a design matrix with an intercept as the wild type genotype, and the coefficients as the FS/+ and EOfAD-like/+ genotypes, and the W_1 coefficient from *RUVg*. Genes were considered differentially expressed (DE) in each specific comparison if the FDR-adjusted p-value was below 0.05.

Enrichment analysis was performed on the KEGG and GO gene sets from the Molecular Signatures Database (MSigDB) [25], with GO terms excluded if the shortest path back to the root node was < 3 steps. We also tested for possible iron dyshomeostasis in the RNA-seq data by performing an enrichment analysis on the iron-responsive element (IRE) gene sets described in [26]. To test for overrepresentation of these gene sets within the DE gene lists, we used *goseq* [27], using the average transcript length per gene to estimate sampling bias for DE genes. For enrichment testing on the entire list of detectable genes, we calculated the harmonic mean p-value [28] of raw p-values from *fry* [29], *camera* [30] and *fgsea* [31, 32]. Gene sets were considered significantly altered if the FDR-adjusted harmonic mean p-value was below 0.05. Visualisation of RNA-seq data analysis was performed using *ggplot2* [33], *pheatmap* [34] and *UpSetR* [35].

### Reproducibility and data availability

The raw fastq files and the gene-level counts have been deposited in the GEO database with accession number GSE158233. All code to reproduce this analysis can be found at https://github.com/UofABioinformaticsHub/20181113_MorganLardelli_mRNASeq.

## Results

We first inspected the transcript-level counts to ensure that expression of the *psen2* alleles was as expected (**Additional File 1**). Consistent with observations in [12], expression of the EOfAD-like transcript was at a similar level to that of the wild type *psen2* transcript in the EOfAD-like/+ brains, while the decreased expression of the FS allele of *psen2* relative to the wild type allele was suggestive of nonsense-mediated decay (NMD). We observed that one of the samples had been incorrectly genotyped during sample preparation and was actually transheterozygous for the EOfAD-like and FS alleles of *psen2*. Therefore, we omitted this sample from subsequent analyses. We also examined the relationship between samples by principal component analysis (PCA) on the gene-level counts. Samples did not cluster by genotype in the PCA plot, supporting that heterozygosity for the *psen2* mutations in this study does not give widespread effects on the brain transcriptome. Additionally, samples did not cluster by sex, consistent with our previous observations that sex does not show extensive effects in zebrafish brain transcriptomes [16, 17, 36]. In the PCA plot, we noticed that one of the wild type brain samples (12_WT_4) separated greatly from the others, appearing to be an outlier (**Additional File 1**). Sample weights, as calculated using the *voomWithQualityWeights* algorithm from the *limma* package [37] on the gene-level counts, revealed that this wild type sample was highly down-weighted. Therefore, we also omitted this sample from subsequent analyses.

### Heterozygosity for an EOfAD-like or an FS mutation in *psen2* results in limited differential expression of genes

To identify which genes were dysregulated due to heterozygosity for the EOfAD-like or FS alleles of *psen2*, we performed differential gene expression analysis using a GLM and likelihood ratio test with *edgeR*. This revealed 32 differentially expressed (DE) genes due to the FS mutation, and 5 DE genes due to the EOfAD-like mutation relative to wild type (**Figure 1, Additional File 2**). Only two genes were seen to be significantly upregulated in both comparisons relative to wild type: *AL929206.1* and *BX004838.1*. However, these genes are currently not annotated and have no known function. We tested for over-representation within the DE genes using *goseq* (using the average transcript length per gene as the predictor variable) of gene ontology (GO) terms which, as the name suggests, use ontologies to annotate gene function; KEGG gene sets which can give insight into changes in various biological pathways and reactions; and our recently defined IRE gene sets [26] which can give insight to possible iron dyshomeostasis. However, due to the low numbers of DE genes in both comparisons, no significantly over-represented GO terms or gene sets were identified. (For the top 10 most significantly over-represented GO terms and gene sets in the DE genes, see **Additional File 3**).

### Heterozygosity for an EOfAD-like or an FS mutation in *psen2* has distinct effects on biological processes in young adult zebrafish brains

We next performed enrichment analysis on the entire list of detectable genes in the RNA-seq experiment to obtain a more complete view on the changes to gene expression due to heterozygosity for the EOfAD-like or FS mutations in *psen2*. Due to the highly overlapping nature of GO terms (many GO terms include many of the same genes), we tested only the KEGG and IRE gene sets. We found statistical evidence for 8 KEGG gene sets in the EOfAD/+ brains and 6 KEGG gene sets in the FS/+ brains to be significantly altered as a group. Notably, no IRE gene sets were found to be significantly altered (**Figure 2A**). Genes in the KEGG_RIBOSOME gene set were significantly altered in both comparisons. However, the genes were mostly downregulated in EOfAD-like/+ brains and upregulated in FS/+ brains (**Figure 2B**). For additional visualisations of our enrichment analysis, see **Additional File 4**.

**Figure 2:**
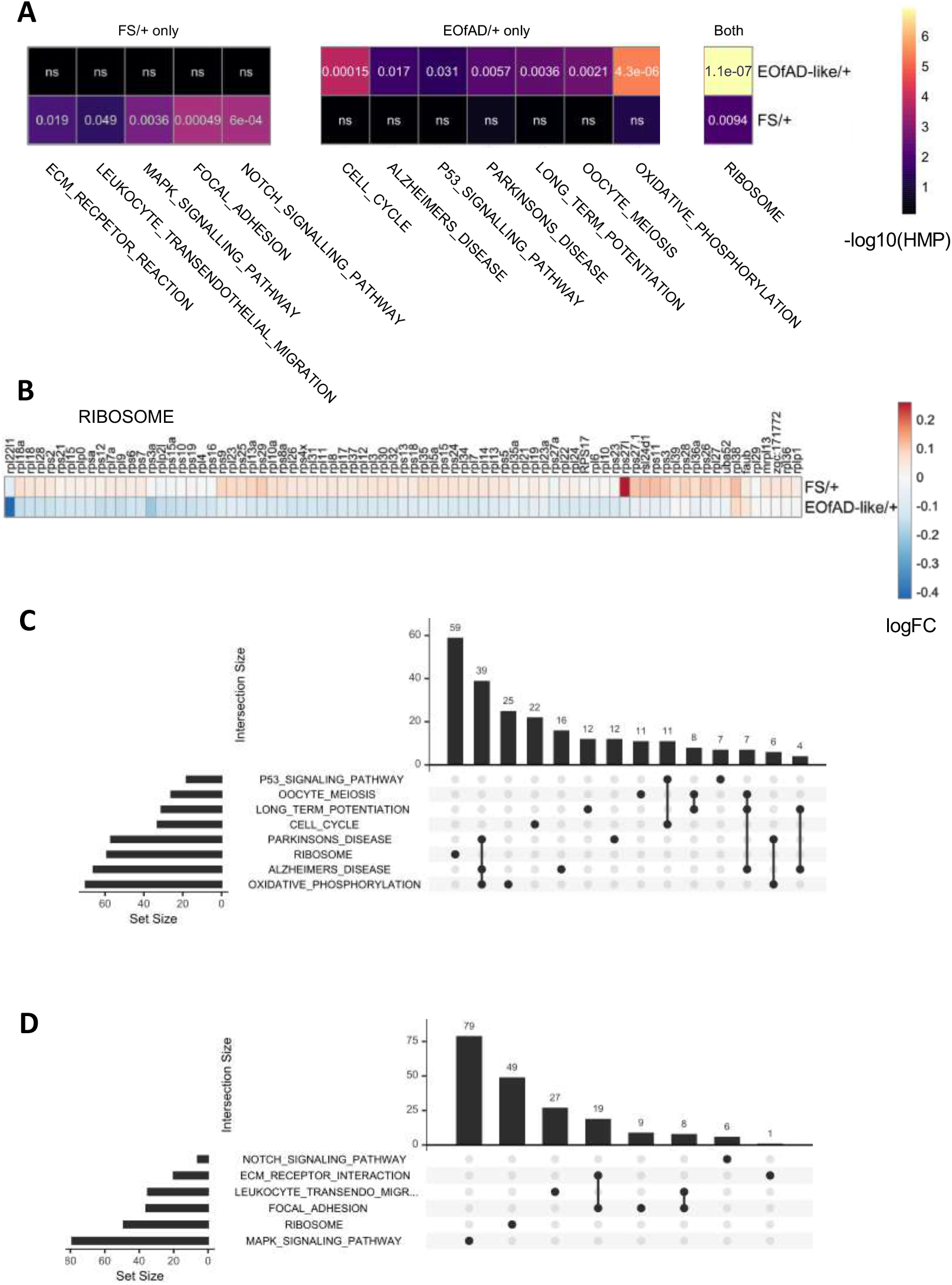
**A)** Heatmap showing the significant KEGG and IRE gene sets in *psen2* heterozygous mutant brains, clustered based on their Euclidean distance. The columns represent the gene sets which are significant in at least one comparison, and the rows are the two comparisons. The colour of each cell represents the significance value with more significant gene sets appearing lighter. The numbers within each cell are the FDR-adjusted harmonic mean p-values (HMP). ns, not significant. **B)** Heatmap indicating the logFC of genes in the KEGG_RIBOSOME gene set in EOfAD/+ and FS/+ brains. **C)** Upset plot showing the overlap of leading edge genes from the *fgsea* algorithm in gene sets found to be altered in EOfAD-like/+ brains, and **D)** FS/+ brains.

To determine whether the statistical significance of the enrichments of the gene sets was being driven by the expression of the same genes, we inspected the genes from the “leading edge” from *fgsea*. (These can be thought of as the core genes driving the enrichment of each gene set). The leading edge genes in each comparison were mostly independent of one another, except for the KEGG gene sets for oxidative phosphorylation, Parkinson’s disease and Alzheimer’s disease, which share 39 genes (**Figure 2C and D**).

## Discussion

Here, we aimed to identify similarities and differences in the effects on brain transcriptomes due to heterozygosity for an EOfAD-like mutation compared to a non-EOfAD-like mutation in *psen2*. To this end we used our BeST strategy, involving analysis of sibling fish raised together in an identical environment. BeST analysis reduces confounding effects in direct comparisons of two heterozygous *psen2* mutations relative to wild type, allowing detection of subtle transcriptome state differences.

We observed subtle, but highly statistically significant, changes to the transcriptomes of the *psen2* mutant fish relative to wild type. At the single gene level, relatively few genes were observed to be statistically significantly DE in each mutant genotype, and differentially enriched biological pathways were not detected in these DE gene lists (**Additional File 3**). As a comparison, heterozygosity for an EOfAD-like mutation in zebrafish *psen1*, (orthologous to the human gene most commonly mutated in EOfAD [38]), gives 251 DE genes in young adult zebrafish brains [15]. However, we were able to identify statistically significant changes in the *psen2* mutant brains by performing enrichment analysis on the entire list of detectable genes in the RNA-seq experiment using an ensemble approach. This involves combining raw p-values by calculation of a harmonic p-value – a strategy thought to be less restrictive than other methods of combining p-values. An additional benefit of this method is that it has been shown specifically to be robust to dependent p-values [28]. The subtlety of the *psen2* mutation effects is consistent with a generally later age of onset, and variable penetrance of EOfAD mutations in *PSEN2* [39–41] relative to EOfAD mutations in *PSEN1* and *APP*.

Interestingly, the pathways which were affected by the two *psen2* mutations showed very little overlap. Gene sets which were significantly altered in EOfAD-like/+ brains were not significantly altered in the FS/+ brains and vice versa. Only one biological process appears to be significantly affected by both mutations: ribosome function (the KEGG_RIBOSOME gene set). However, in general, the direction of differential expression of genes in this gene set was observed to be opposite in the two mutants, implying different mechanisms of action of each mutation (**Figure 2B**). The low concordance between the brain transcriptome effects of these two mutations supports that presenilin EOfAD mutations do not act through a simple loss of function but, instead, through a gain of function mechanism connected with encoding an abnormal, but “full-length” protein [13]. Notably, the EOfAD-like mutation of *psen2* is predicted to affect the process of long term potentiation thought to be critical for memory formation. We also observed the dysregulation of genes involved in controlling the cell cycle. Genes driving the enrichment of this gene set (i.e. the leading edge genes) include the *minichromosome maintenance* (*mcm*) genes (see **Additional File 4**), which we observed to be significantly dysregulated due to heterozygosity for an EOfAD-like mutation in *psen1* in 7 day old zebrafish larvae [42]. Together, these results provide support for the “two-hit hypothesis” of Mark Smith and colleagues [43], which postulates that “cell cycle events” (CCEs) involving inappropriate attempts at cell division by neurons are critical for development of AD. The second “hit” of this hypothesis is oxidative stress, which is an anticipated outcome of disturbance of oxidative phosphorylation. Notably, the EOfAD-like mutation of *psen2* is predicted to affect oxidative phosphorylation, which is in common with all the other EOfAD-like mutations of genes we have examined in zebrafish [16, 18, 26, 44].

One of the most characterised functions of presenilin proteins is their role as the catalytic core of gamma-secretase (*γ*-secretase) complexes. *γ*-secretase activity is responsible for cleavage of a wide range of type I transmembrane proteins including AβPP and NOTCH. We observed that genes involved in the Notch signalling pathway were only significantly altered in FS/+ genotype brains, indicating that *γ*-secretase activity may be altered by the FS mutation. Interestingly, inspection of the direction of changes to gene expression in this pathway’s gene set suggest an increase in *γ*-secretase activity (**Additional File 4**), as *notch, delta* and, particularly, *hes* genes (that are Notch signalling targets) are all observed as upregulated. This seems counterintuitive, as the FS mutation results in a truncated protein lacking critical aspartate residues needed for *γ*-secretase activity (**Figure 1B**). A minor, (although statistically non-significant) increase in expression of *psen1* is observed in FS/+ brains (also in EOfAD-like/+ brains) (**Additional File 4**) raising the possibility that transcriptional adaptation (formerly known as “genetic compensation”) might explain an apparent increase in *γ*-secretase activity [45]. Since presenilin holoproteins are known to form multimeric complexes, we have suggested previously that this may be involved in the regulation of the conversion of the holoprotein into its *γ*-secretase-active form by endoproteolysis [13]. N-terminal fragments of presenilin proteins are also known to multimerise [46] and the possibility exists that these may disrupt holoprotein multimer formation/stability. In our previous publication identifying the EOfAD-like and FS mutations of *psen2* analysed here, we saw that both mutations reduced adult melanotic surface pigmentation when homozygous [12].

This implied that both mutations reduce *γ*-secretase activity, at least in melanosomes. These apparently conflicting observations support that analysis of brain biochemistry in these EOfAD-like and FS zebrafish *psen2* mutants may reveal important details of the relationship between presenilin holoproteins and their *γ-* secretase-active endoproteolysed forms including between the holoproteins encoded by *PSEN1* and *PSEN2*.

### Conclusion

In conclusion, our results support that frameshift and reading frame-preserving mutations in the *presenilins* have distinct effects on the brain transcriptome, consistent with the reading frame preservation rule of presenilin EOfAD-causative mutations. The data presented here, along with our growing collection of BeST analyses, indicates that changes to mitochondrial and ribosomal functions are effects-in-common of heterozygosity for EOfAD-like mutations in different genes. These may be early cellular stresses which eventually lead to AD pathology, and warrant investigation for discovery of therapeutic targets.

## Declarations

The authors have no conflict of interest to declare.

## Acknowledgements

The authors would like thank FBRI LLC for use of the presenilin protein schematic in Figure 1. This work was supported with supercomputing resources provided by the Phoenix HPC service at the University of Adelaide, and by grants GNT1061006 and GNT1126422 from the National Health and Medical Research Council of Australia (NHMRC). KB was supported by an Australian Government Research Training Program Scholarship. HJ was supported by an Adelaide Scholarship International scholarship from the University of Adelaide.

## Additional Files

**Additional File 1.**
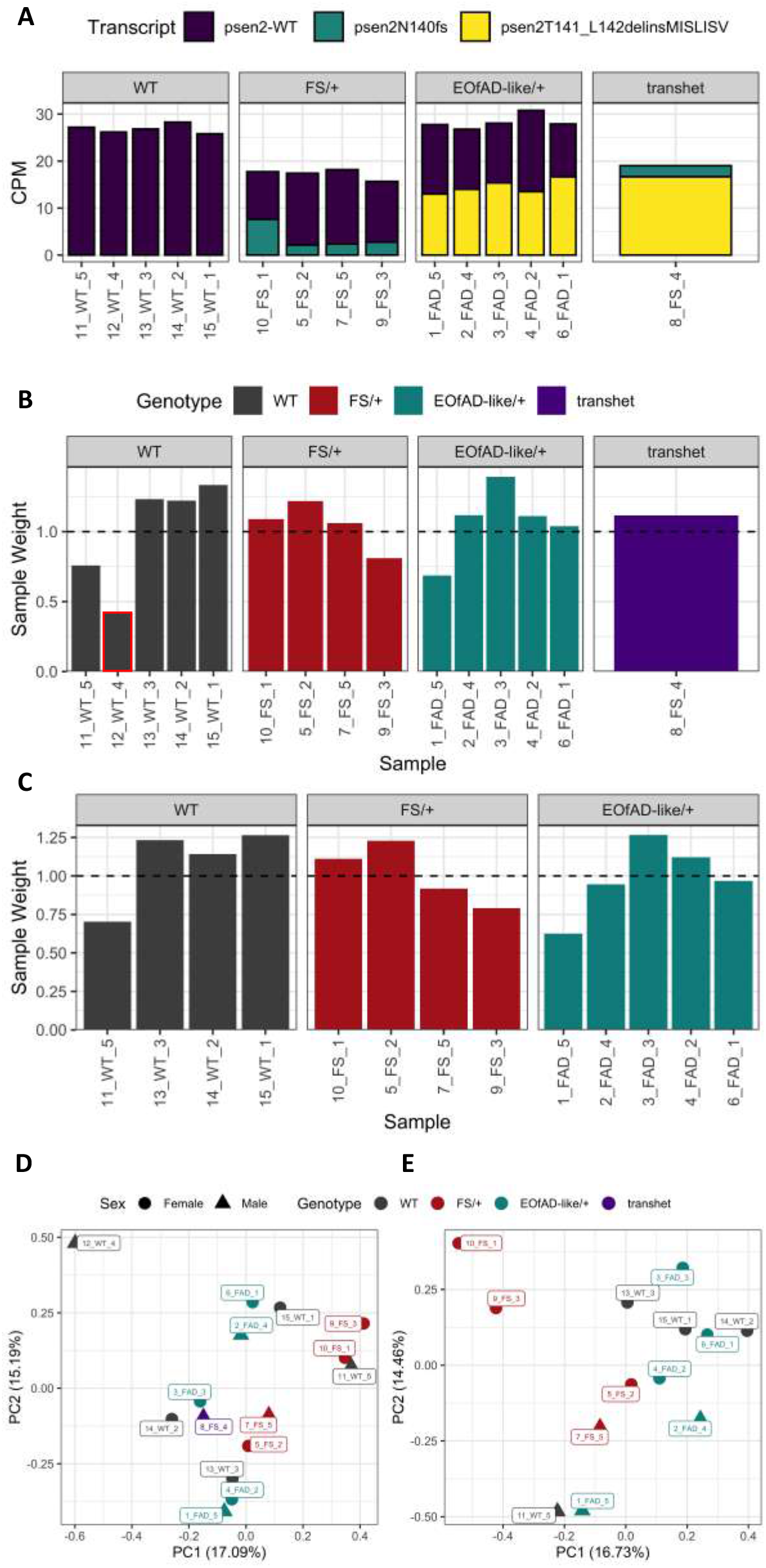
**A.** Allele-specific expression (in counts per million, CPM) of *psen2* transcripts in young adult zebrafish brains. Reduced expression of the *psen2^N140fs^* allele is observed in FS/+ brains, consistent with our previous observation that transcripts of this allele are subject to nonsense-mediated decay. Sample 8_FS_4 does not express the wild type allele of *psen2* and is actually a transheterozygous (transhet) sample. Therefore, it was omitted from the analysis. **B.** Sample weights as calculated by the *voomWithQualityWeights* algorithm on all samples sequenced. Sample 12_WT_4 is highly downweighted relative to all other samples and was omitted from subsequent analysis. **C.** Sample weights recalculated after exclusion of samples 8_FS_4 and 12_WT_4. **D.** Principal component 1 (PC1) against PC2 from a principal component analysis (PCA) on all samples of the experiment. Sample 12_WT_4 does not cluster with the other samples. **E.** PCA plot of the *RUVseq*-normalised counts after exclusion of samples 8_FS_4 and 12_WT_4.

Additional File 2: Full results of differential gene expression analysis.

Additional File 3: Over-representation analysis using goseq.

**Table 1:** The KEGG, GO and IRE gene sets approaching being over-represented in the DE genes list for the FS mutation. The 5 DE genes due to the EOfAD-like mutations are not found in any of the gene sets. Therefore, the results of the overrepresentation analysis are not shown.

| Gene set | p-value | Number of DE genes in gene set | Number of genes in gene set | FDR adjusted p-value |
| --- | --- | --- | --- | --- |
| GO_PHOSPHOTRANSFERASE_ACTIVITY_NITROGENOUS_GROUP_AS_ACCEPTOR | 1.94E-05 | 2 | 5 | 0.28952 |
| GO_TRNA_3_END_PROCESSING | 6.94E-05 | 2 | 9 | 0.517902 |
| GO_NUCLEOSIDE_DIPHOSPHATE_METABOLIC_PROCESS | 1.01E-03 | 3 | 142 | 1 |
| KEGG_ALZHEIMERS_DISEASE | 1.11E-03 | 3 | 146 | 1 |
| GO_NCRNA_3_END_PROCESSING | 1.39E-03 | 2 | 39 | 1 |
| GO_PYRIMIDINE_NUCLEOBASE_TRANSPORT | 2.85E-03 | 1 | 2 | 1 |
| GO_UREA_TRANSMEMBRANE_TRANSPORTER_ACTIVITY | 2.85E-03 | 1 | 2 | 1 |
| GO_TRNA_BINDING | 3.58E-03 | 2 | 63 | 1 |
| GO_GLYCEROL_CHANNEL_ACTIVITY | 4.26E-03 | 1 | 3 | 1 |
| GO_UREA_TRANSPORT | 4.27E-03 | 1 | 3 | 1 |

**Additional File 4:**
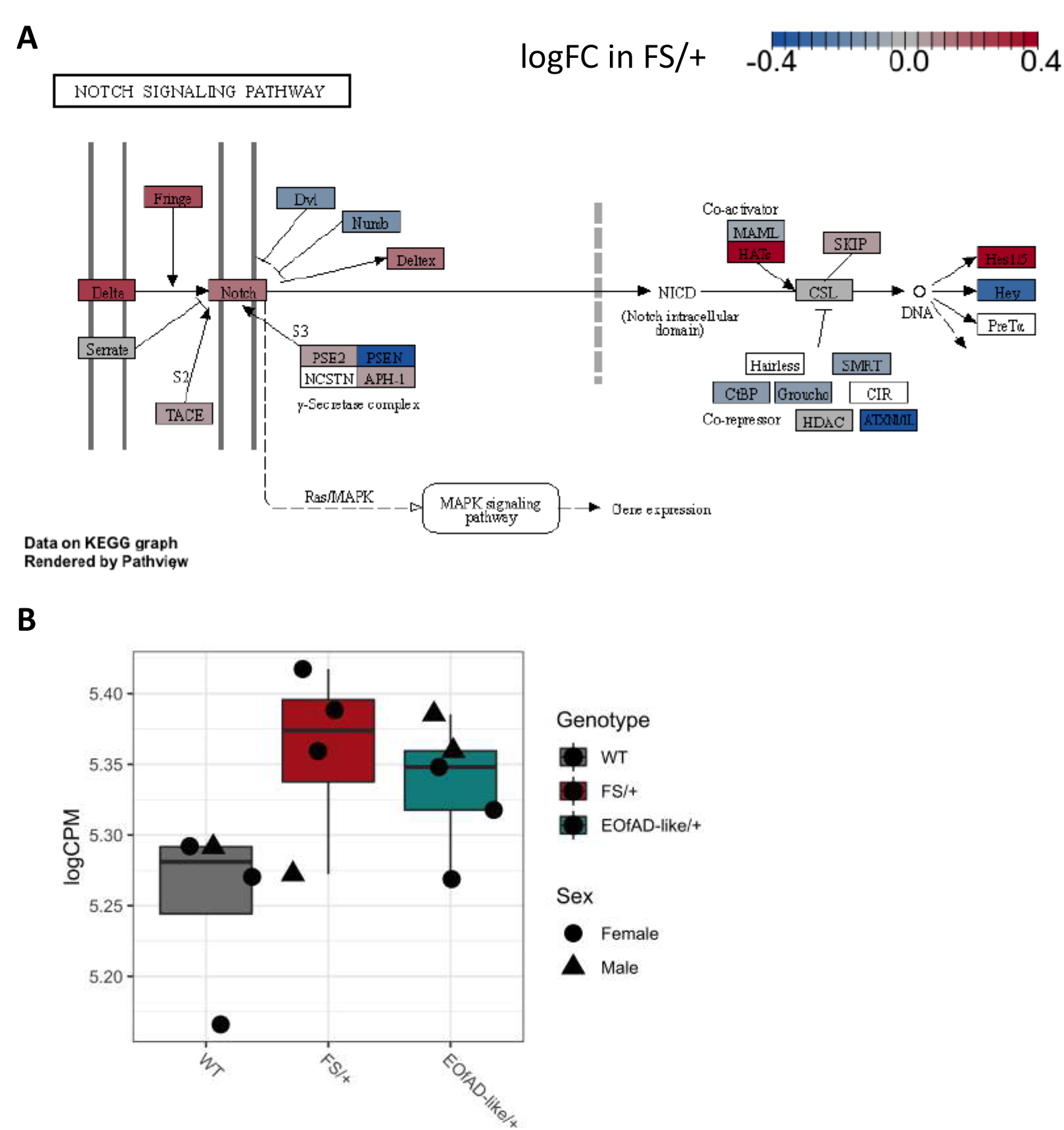
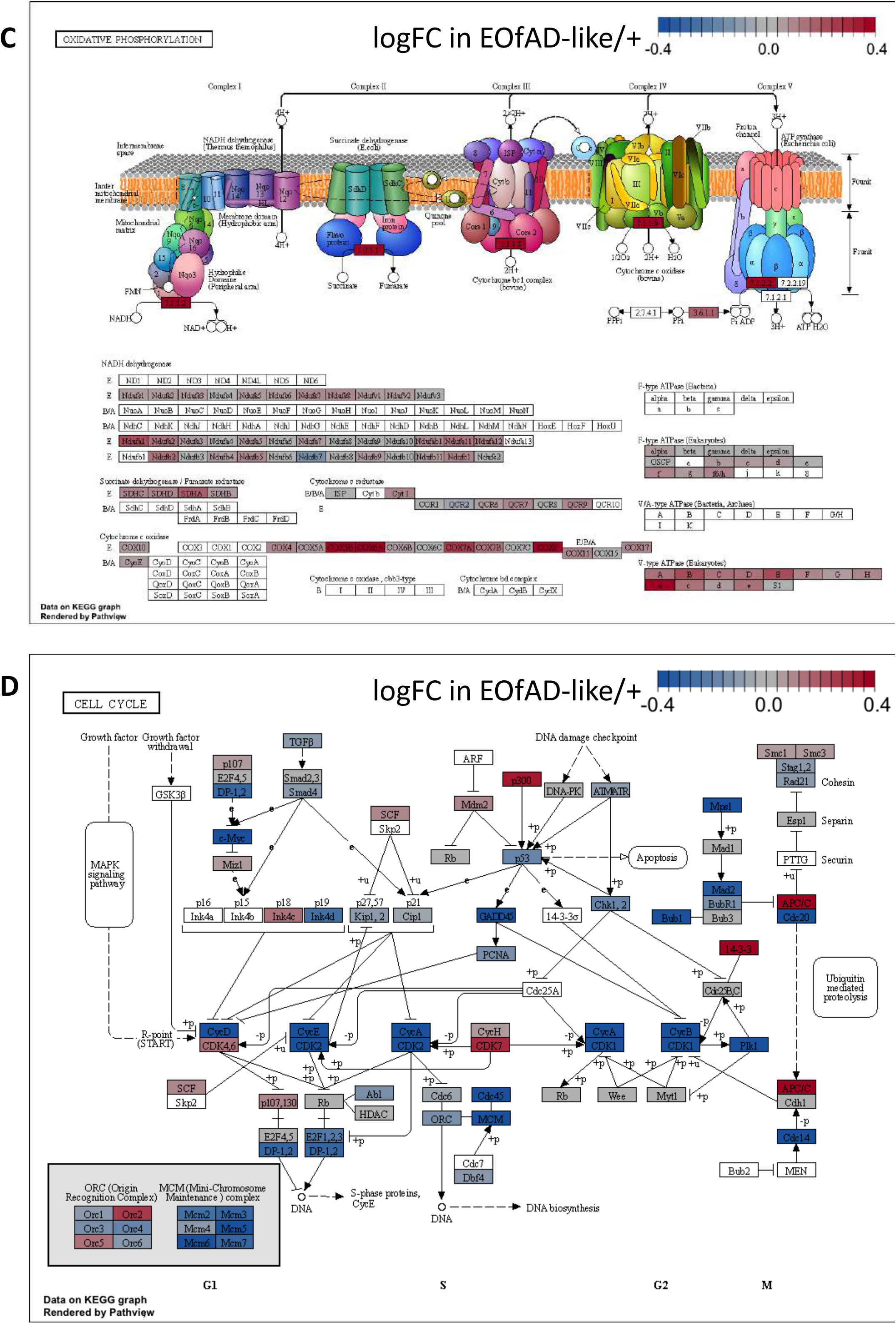
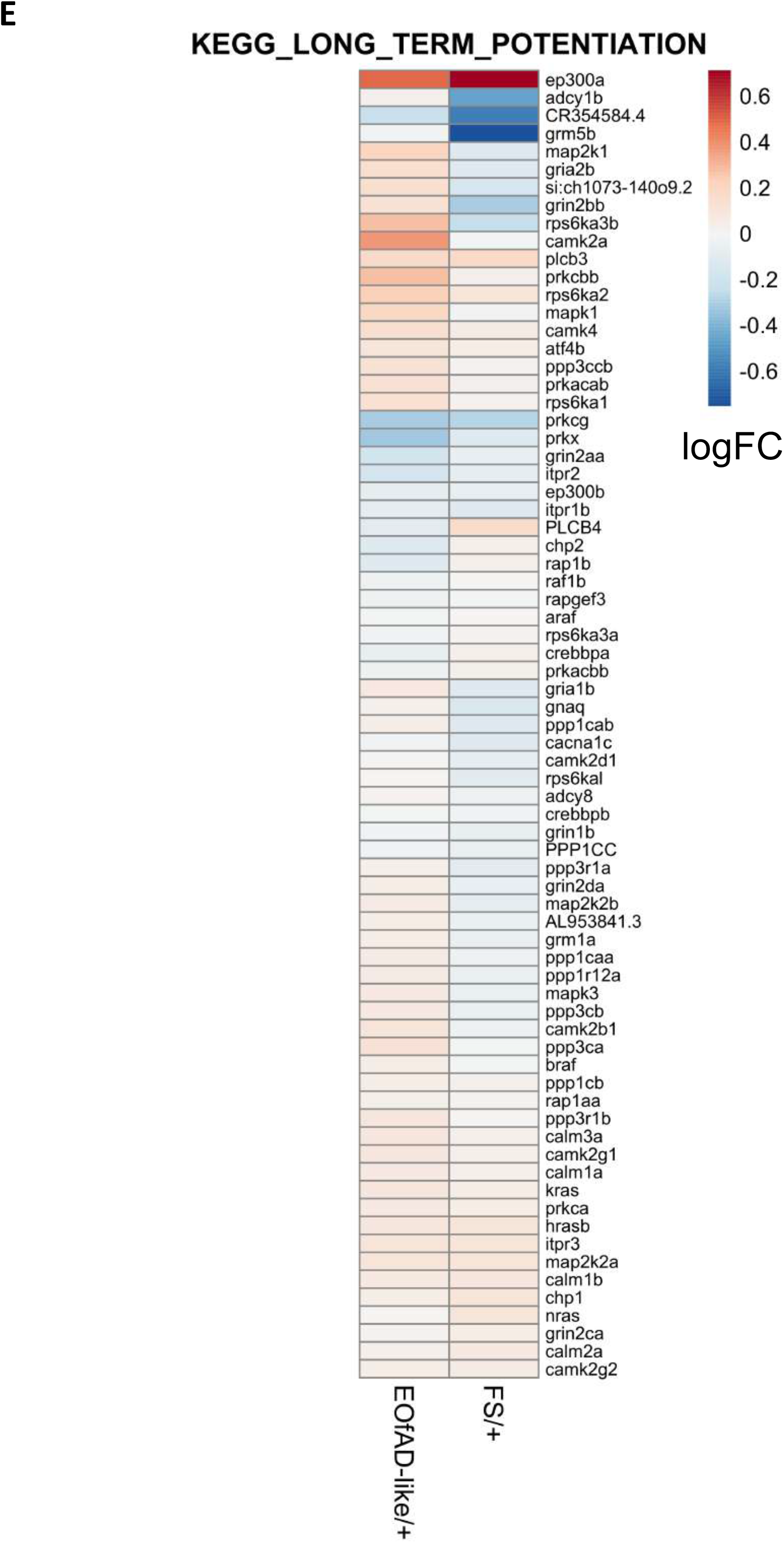
Additional RNA-seq visualisations. **A.** Pathview [47] visualisation of the logFC of genes in the KEGG_NOTCH_SIGNALLING_PATHWAY gene set in FS/+ brains. **B.** Expression of *psen1* in log counts per million (logCPM). **C.** Pathview [47] visualisation of the logFC of genes in the KEGG_OXIDATIVE_PHOSPHORYLATION gene set in EOfAD-like/+ brains. **D.** Pathview [47] visualisation of the logFC of genes in the KEGG_CELL_CYCLE gene set in EOfAD-like/+ brains. **E.** Heatmap of the logFC of genes in the KEGG_LONG_TERM_POTENTIATION gene set in EOfAD-like/+ brains.

